# The porin MspA is a phage-specific determinant of mycobacteriophage infection

**DOI:** 10.64898/2026.10.08.757660

**Authors:** Doyeon Kim, Mickaël Maziero, Kathryn A. Eckartt, Michael A. DeJesus, Grisna Isabel Prensa, Andrea García Navarro, David R. Parker, Pierre Santucci, Jeremy M. Rock, Andrea Fossati

**Affiliations:** Department of Microbiology, Tumor and Cell Biology (MTC), Karolinska Institutet, Biomedicum C9, Solnavägen 9, Solna, 17165, Sweden; Science for Life Laboratory, Karolinska Institutet, Tomtebodavägen 23A, Solna, 17165, Sweden; Laboratoire d’Ingénierie des Systèmes Macromoléculaires (LISM), IM2B, Aix Marseille Univ, CNRS, 31 Chemin Joseph Aiguier, Marseille, 13402, France; Laboratory of Host-Pathogen Biology, The Rockefeller University, 1230 York Avenue, New York, NY, 10065, USA

## Abstract

Mycobacteriophages are being developed as therapeutics for mycobacterial infections, but their highly variable host ranges limit this application and the bacterial factors that determine whether infection succeeds remain poorly defined. The determinants characterized so far are dominated by cell envelope lipids and glycolipids. Here, we performed a genome-wide CRISPRi screen in *Mycobacterium smegmatis* to identify host functions associated with susceptibility to BPs**Δ**33HTH HRM10. After prolonged phage challenge, the outer membrane porin genes *mspA* and *mspC* were among the most strongly enriched hits, whereas the closely related paralogs *mspB* and *mspD* were not enriched despite sharing high amino-acid identity across the family. Deletion of *mspA* or *mspC* reduced BPs**Δ**33HTH HRM10 plaque formation and produced smaller, more turbid plaques, with the stronger phenotype following loss of *mspA*, the major outer membrane porin of *M. smegmatis*. Across a panel of 25 mycobacteriophages, 11 showed either a more than tenfold reduction in titer or loss of discrete plaque formation on **Δ***mspA*, while the remainder were largely unaffected. These findings identify outer membrane porins as phage-dependent determinants of mycobacterial susceptibility, revealing an additional layer of specificity in phage infection beyond previously characterized cell envelope lipids and glycolipids.

## Introduction

Bacteriophage therapy is increasingly being investigated for the treatment of mycobacterial infections, including those caused by *Mycobacterium abscessus*[1–4]. Its broader therapeutic application, however, is constrained by the limited and highly variable host ranges of individual mycobacteriophages[5]. In a recent screening effort involving isolates from 200 culture-positive patients with symptomatic mycobacterial disease, at least one active lytic phage was identified for only 55 isolates[3]. Substantial variation in phage susceptibility has likewise been observed among *M. abscessus* clinical isolates[1]. Despite the remarkable diversity of mycobacteriophages, the bacterial factors that determine whether productive infection succeeds or fails remain poorly defined[6]. Studies of mycobacteriophage susceptibility have identified surface glycolipids as important host determinants[7, 8]. Trehalose polyphleates (TPPs) support efficient adsorption and infection by BPs and Muddy in *M. abscessus* and *M. smegmatis*[9]. More recently, loss of the mycolyltransferase Ag85A was shown to reduce BPs adsorption and infection through surface-lipid remodeling, with excess trehalose dimycolate proposed to restrict access to TPPs[10]. Together, these studies show that mycobacteriophage adsorption can be disrupted either through alteration or loss of a phage interacting surface glycolipid or through remodeling of the surrounding lipid environment that changes access to that surface determinant[7, 9, 10]. Thus, the experimentally characterized host determinants of mycobacteriophage adsorption have so far been dominated by lipid and glycolipid components of the cell envelope, whereas the contribution of other classes of bacterial factors remains much less well defined. A systematic search for such host determinants is limited by the genetic approaches traditionally used to study mycobacteriophage susceptibility. Spontaneous phage resistant mutants, targeted gene disruption and transposon mutagenesis have provided important insights into mechanisms of phage resistance and host dependence[1, 7, 9]. However, these approaches generally rely on the recovery of viable loss-of-function mutants and therefore provide limited access to essential genes or genes whose complete loss causes severe fitness defects. Indeed, transposon based analyses identified 625 genes or essential domains required for optimal *in vitro* growth in *M. tuberculosis* and 403 genes classified as essential or growth defective in *M. smegmatis*[11, 12], making many of these genes difficult to interrogate directly for their contribution to phage susceptibility using conventional loss of function screens. CRISPR interference (CRISPRi) provides a complementary approach by enabling tunable repression of gene expression without requiring complete gene inactivation[13]. Genome-scale CRISPRi libraries consequently extend functional genetic screening to genes that are difficult to access using conventional loss of function approaches[14, 15]. Here, we used genome-wide CRISPRi screening in *M. smegmatis* to identify bacterial factors influencing mycobacteriophage susceptibility. The screen identified the outer membrane porins MspA and MspC as candidate host factors, whose contributions to infection were subsequently validated through targeted genetic perturbations. Screening a panel of 25 mycobacteriophages revealed substantial variation in porin dependence, including differences between parental phages and derivatives selected for expanded infectivity toward *M. abscessus*. Together, these findings establish outer membrane proteins as an additional class of determinants of mycobacteriophage susceptibility and demonstrate that their contribution depends on the infecting phage.

## Results

### Establishment of a genome-wide CRISPRi screen for mycobacteriophage susceptibility

A pooled CRISPRi screen was designed to identify bacterial genes that influence mycobacteriophage susceptibility in *M. smegmatis*. The RLC11 library, comprising 168,865 sgRNAs targeting genes across the *M. smegmatis* genome together with 7,421 non-targeting controls, was induced with anhydrotetracycline (aTc) and subsequently challenged with phage (Fig. 1A). Under phage selection, enrichment of targeting sgRNAs indicates that reduced expression of the corresponding gene provides a relative fitness advantage, whereas depletion indicates reduced bacterial fitness during phage exposure. To establish a screening condition that provided strong and sustained selective pressure, we compared the growth of *M. smegmatis* following infection with Muddy and BPsΔ33HTH HRM10, which were selected from a panel of mycobacteriophages previously used in the treatment of clinical *M. abscessus* infections. BPsΔ33HTH HRM10 produced stronger and more sustained growth inhibition than Muddy and was therefore selected for further optimization of the screening conditions. Subsequent testing across multiplicities of infection (MOI) from 0.01 to 10 showed pronounced growth inhibition at MOI 1 and above. As MOI 1 was the lowest condition that produced a strong inhibitory phenotype, it was selected for the genome-wide CRISPRi screen (Fig. 1B,C). During the early phase of BPsΔ33HTH HRM10 infection, cultures showed pronounced growth suppression, whereas partial regrowth of the bacterial population was observed following prolonged phage exposure. We therefore selected 18 h, when phage mediated growth inhibition was prominent, and 96 h, when substantial regrowth was apparent, as complementary sampling points to examine host gene effects during early and prolonged phage selection (Fig. 1C). Before proceeding to gene-level analysis, we examined sgRNA abundance across the experimental time course. Relative to the starting library, targeting sgRNAs showed an overall shift toward negative log_2_ fold change (L2FC) values, with a distribution mean of *−*1.523 at 18 h and *−*1.504 at 96 h and 91.7% and 91.0% of sgRNAs falling below zero, consistent with depletion of knockdowns associated with reduced bacterial fitness. Non-targeting controls remained centered close to zero at both times (mean *−*0.016 and *−*0.020) (Fig. 1D,E). To distinguish these general fitness associated changes from phage specific selection, we directly compared phage treated cultures with their time matched no-phage controls. At both 18 h and 96 h, sgRNAs targeting essential and nonessential genes spanned a broader range of positive and negative L2FC values than non-targeting controls, indicating differential selection of individual knockdowns during phage exposure (Fig. 1F). These results established that the screen captured differential sgRNA selection during phage exposure, providing the basis for subsequent gene-level analysis.

**Fig. 1.**
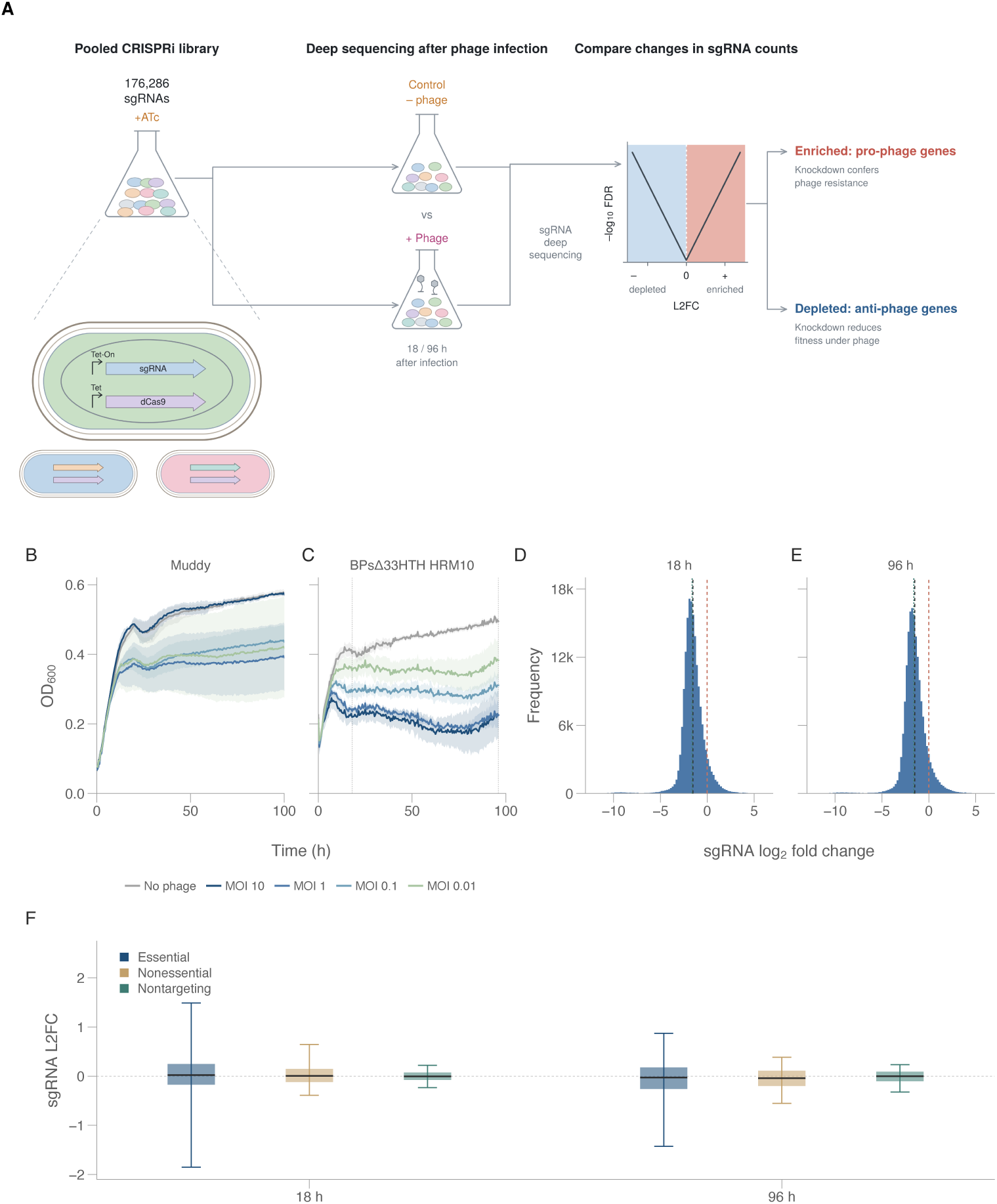
Experimental design and sgRNA abundance distributions in the CRISPRi phage screen. **A.** Pooled CRISPRi screen in *Mycobacterium smegmatis* mc^2^155: induction of the RLC11 library with anhydrotetracycline (aTc), division into phage-exposed and no-phage arms, and deep sequencing of sgRNA abundance. **B, C.** Growth of *M. smegmatis* mc^2^155 after exposure to Muddy (**B**) or BPsΔ33HTH HRM10 (**C**) at the indicated multiplicities of infection (MOIs); uninfected cultures received an equal volume of SM buffer. Lines and shading, mean and sample standard deviation of three recorded OD_600_ curves. Dotted lines in **C**, the 18 h and 96 h sampling times. **D, E.** sgRNA log_2_ fold change relative to the original library at 18 h (**D**) and 96 h (**E**), for all 176,286 sgRNAs. Vertical lines mark zero, the distribution mean and median, and the mean of the non-targeting controls. **F.** Phage-associated sgRNA L2FC at MOI 1 relative to the time-matched no-phage control, at 18 h and 96 h, by guide class (55,098 essential, 113,767 nonessential and 7,421 non-targeting sgRNAs per time point). Boxes, interquartile range; central line, median; whiskers, 5^th^ to 95th percentile.

### Genome-wide CRISPRi screening identifies MspA and MspC as candidate mycobacteriophage host factors

Gene-level analysis identified multiple genes whose targeting sgRNAs were enriched or depleted following BPsΔ33HTH HRM10 exposure. At 96 h, the outer membrane porin genes *mspA* (MSMEG_0965, L2FC = +2.25) and *mspC* (MSMEG_5483, L2FC = +2.98) were among the prominent enriched hits, indicating that reduced expression of either gene was associated with a relative fitness advantage during phage challenge (Fig. 2A,B). At the same time point, several prominent depleted hits were associated with cell envelope functions, including the MmpL-family transporter MmpL3 (MSMEG_0250, L2FC = *−*1.98), the LppX/LprAFG-family lipoprotein MSMEG_ 3070 (L2FC = *−*3.24), and LpqB (MSMEG_1876, L2FC = *−*3.83) (Fig. 2B). Functional classification revealed broader directional patterns across the screen. At 18 h, cell wall associated genes formed the largest displayed category among depleted hits, accounting for 17.9% of the total. In contrast, tRNA related genes and tRNA lig-ases were prominent among enriched hits, together accounting for 20.3% of enriched hits at 18 h and 31.9% at 96 h (Fig. 2D,E). Thus, cell wall associated knockdowns were preferentially represented among depleted hits, particularly at 18 h, whereas tRNA related knockdowns were consistently represented among enriched hits at both time points. Comparing gene-level effects directly between the two sampling times showed that most genes changed little, while the depleted envelope associated candidates became more strongly negative and *mspA* and *mspC* moved furthest toward enrichment at 96 h (Fig. 2C). Because the screen was sampled at both 18 h and 96 h, we next used MAGeCK maximum-likelihood estimation to assess changes in phage associated effects over time. The analysis revealed heterogeneous temporal responses among the screen candidates. For *mspA*, the phage-associated coefficient (*β*, the MLE effect size on a log fold change scale, where the sign gives the direction of selection and the magnitude its strength) increased from *β* = 0.599 at 18 h to *β* = 1.562 at 96 h, while that of *mspC* increased from *β* = 0.236 to *β* = 0.869. Neither trend reached the predefined threshold for a significant time-dependent phage effect (time*×*phage interaction q = 0.057 for *mspA* and q = 0.153 for *mspC*), and both are therefore reported as descriptive estimates (Fig. 3A). We therefore examined the primary screen evidence for the Msp porins in greater detail. Guide-level analysis showed that multiple sgRNAs targeting distinct positions within *mspA* and *mspC* displayed predominantly positive phage-associated effects, supporting the corresponding gene-level enrichment (Fig. 3B). To determine whether this enrichment extended to other members of the Msp porin family, the four Msp paralogs were compared at both the sequence and phage-associated effect levels. All four carry the same domain organization, a cleaved N-terminal signal peptide followed by a single Pfam porin domain (PF09203) that spans essentially the entire mature chain, shown for MspA and MspC in Fig. 3C, and their signal-peptide-trimmed mature proteins share 89.7–98.9% pairwise amino-acid identity (Fig. 3D). Despite this high degree of sequence conservation, the sgRNAs targeting *mspB* and *mspD* showed substantially weaker effects at 96 h (Fig. 3E), with gene-level L2FC values of +0.14 and +0.36, respectively, and neither gene met the enrichment criteria. Thus, the strong phage-associated enrichment of *mspA* and *mspC* was not a general feature of the Msp porin family, supporting their selection for targeted genetic validation.

**Fig. 2.**
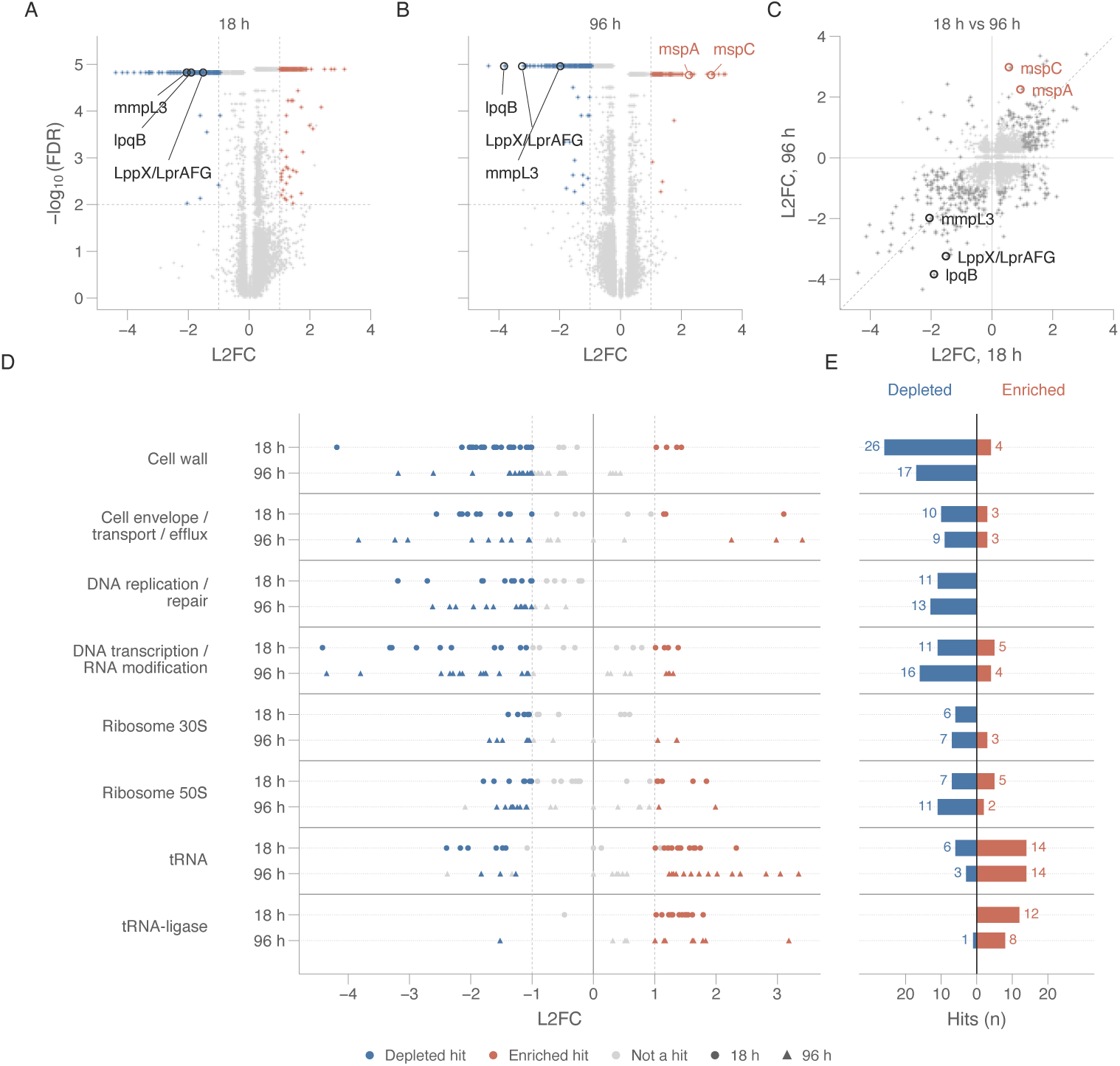
Genome-wide CRISPRi profiling of phage-associated gene effects and their functional distribution. **A, B.** Gene-level effects at 18 h (**A**) and 96 h (**B**) after phage exposure at MOI 1, relative to the time-matched no-phage control. Each point is one gene, shown as L2FC against *−*log_10_(FDR) for the direction with the lower FDR. Red, enriched; blue, depleted (FDR < 0.01, |L2FC| *≥* 1 and at least two supporting sgRNAs); grey, below criteria. Dashed lines, the FDR and L2FC thresholds. Genes sharing the smallest FDR returned by MAGeCK form the band at the top of each panel. Labelled genes are circled; *mspA* and *mspC* are labelled in **B**. **C.** Gene-level L2FC at 18 h against 96 h. Dark grey, genes meeting the hit criteria at either time point; dashed line, *y* = *x*. **D.** Hits by functional group, with one row per time point and values at 18 h (circles) and 96 h (triangles) on a shared L2FC axis. Blue, depleted hit; red, enriched hit; grey, below the hit criteria. Solid rules separate the functional groups and dotted rules mark each row. **E.** Number of hits per functional group in each direction and time point, diverging about zero: blue to the left, depleted; red to the right, enriched. Rows and rules as in **D**. Totals across all groups are 145 depleted and 128 enriched hits at 18 h, and 174 and 69 at 96 h; hits classified as Other are counted in those totals but are not displayed.

**Fig. 3.**
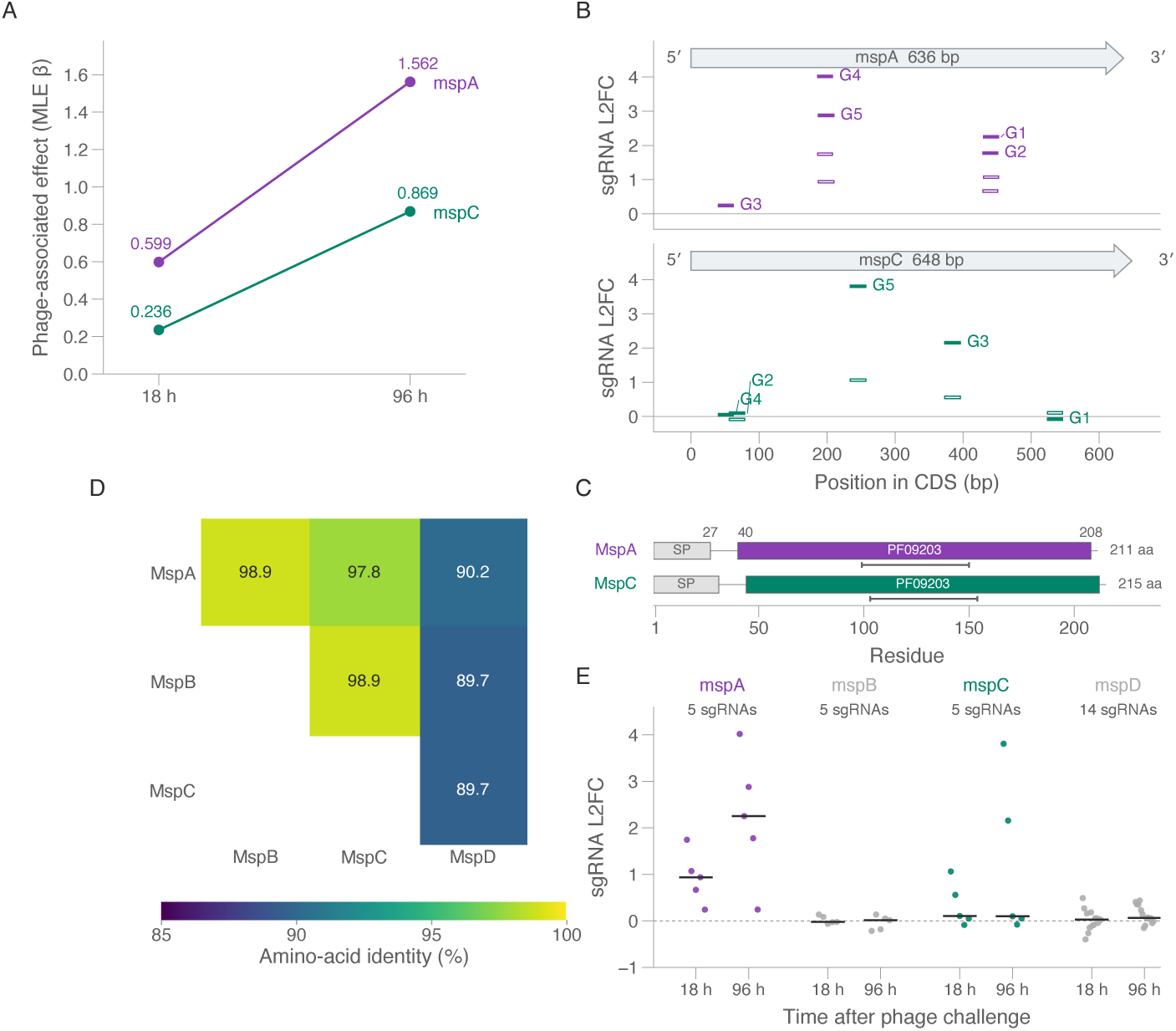
Comparative analysis of phage-associated responses across the Msp porin family. **A.** MAGeCK maximum-likelihood estimate of the phage-associated coefficient (*β*) for *mspA* (MSMEG_0965, purple) and *mspC* (MSMEG_5483, green) at 18 h and 96 h, relative to the time-matched no-phage control. Labels give *β* to three decimal places. **B.** sgRNA L2FC along the *mspA* (top) and *mspC* (bottom) coding sequences. Bar width is the sgRNA target interval, not an uncertainty; open bars, 18 h; filled bars, 96 h. G1–G5 identify individual sgRNAs and the arrow gives coding sequence orientation and length. **C.** Domain architecture of MspA and MspC on a shared residue axis: the cleaved signal peptide (SP, grey), the Pfam porin domain PF09203, and, as a rail beneath each track, the central ribbon domain. Residue boundaries are numbered on the MspA track; precursor lengths are given at the right of each track. Signal-peptide boundaries are those recorded for the supplied precursor and cleaved sequences; domain boundaries are the InterPro assignments for UniProt A0QR29 (MspA) and A0R3I3 (MspC). **D.** Pairwise amino-acid identity among the mature MspA, MspB, MspC and MspD proteins, with signal peptides removed before alignment. The six unique comparisons are shown as an upper triangle. **E.** sgRNA L2FC for *mspA–D* at 18 h and 96 h; horizontal lines, medians. The number of sgRNAs targeting each gene is given above each pair.

### Genetic perturbation of MspA and MspC alters susceptibility to BPsΔ33HTH HRM10

To validate the *mspA* and *mspC* hits identified in the pooled CRISPRi screen, *M. smegmatis* Δ*mspA* and Δ*mspC* deletion mutants were generated and compared with WT. The Δ*mspA* and Δ*mspC* genotypes were verified by locus specific PCR (Supplementary Fig. S1C). In the absence of phage, both deletion mutants showed growth broadly comparable to WT, with only modest differences in their growth trajectories and stationary phase OD_600_ (Fig. 4A). Whole cell proteomics was additionally used to assess changes in Msp porin abundance in the deletion strains. Because of the high sequence similarity among Msp paralogs, individual proteins could not be quantified unambiguously. At the protein-group level, the shared MspA/MspB/MspC signal was approximately 0.25-fold that of WT in Δ*mspA* (q = 0.05005) and 0.68-fold in Δ*mspC* (q = 0.805) (Fig. 4B). This pattern is consistent with previous studies identifying MspA as the major porin of *M. smegmatis*, with *mspA* deletion producing a substantially greater reduction in total porin abundance than deletion of other Msp paralogs[16]. We next examined whether loss of either porin altered susceptibility to BPsΔ33HTH HRM10. In liquid culture, phage exposure caused strong growth suppression of WT, whereas growth inhibition was markedly attenuated in both Δ*mspA* and Δ*mspC*. Both deletion strains continued to increase in optical density under conditions in which WT cultures showed little net growth (Fig. 4C). In spotting assays, BPsΔ33HTH HRM10 formed plaques on both deletion strains, but the plaques were smaller and more turbid than those formed on WT, with a more pronounced phenotype on Δ*mspA* (Fig. 4D). To determine whether partial repression of the same genes produced a similar phenotype, the two sgRNAs showing the strongest positive phage-associated L2FC values in the pooled screen were selected for each gene. Independent CRISPRi strains carrying each selected sgRNA were then tested under the same spotting conditions. For BPsΔ33HTH HRM10, these strains showed more modest plaque phenotypes than the corresponding deletion mutants (Fig. 4D). Taken together, these results indicate that both MspA and MspC contribute to susceptibility to BPsΔ33HTH HRM10, with loss of either porin reducing phage-mediated growth inhibition and altering plaque formation, and with a stronger overall phenotype following *mspA* deletion.

**Fig. 4.**
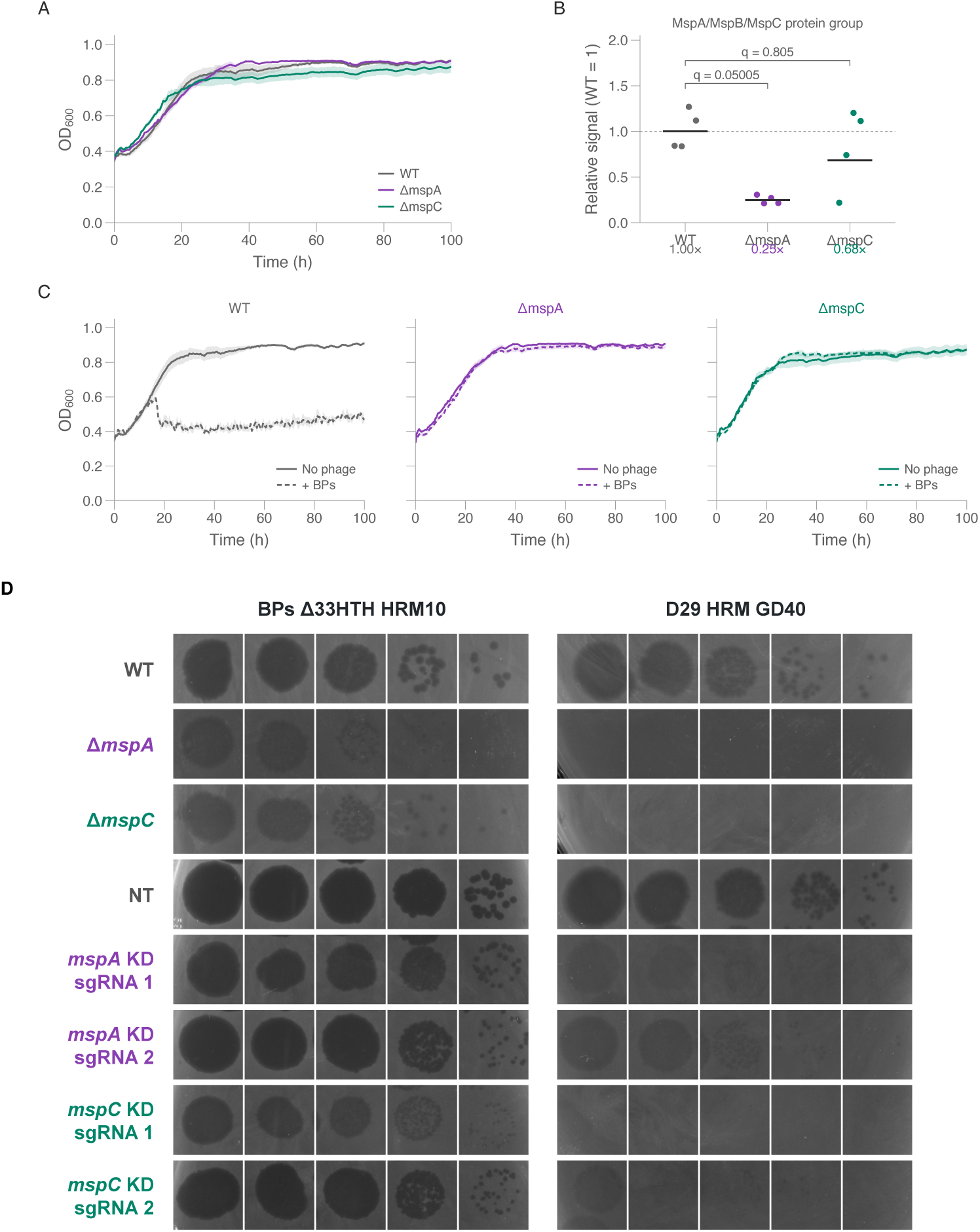
Genetic validation of MspA and MspC as phage susceptibility factors. **A.** Growth of WT, Δ*mspA* and Δ*mspC M. smegmatis* in the absence of phage, monitored by OD_600_. Lines and shading, mean and sample standard deviation of three recorded curves. **B.** Whole-cell abundance of the shared MspA/MspB/MspC protein group, which the detected peptides do not resolve into individual paralogs. Points, four biological replicates; horizontal lines, geometric means normalized to the WT geometric mean; dashed line, the WT level. *q* values are Benjamini–Hochberg-adjusted two-sided Welch tests on normalized log_2_ intensities. **C.** Growth of each strain in the presence (dashed) or absence (solid) of BPsΔ33HTH HRM10, one panel per strain. Lines and shading as in **A**. **D.** Spot assays of BPsΔ33HTH HRM10 and D29 HRMGD40 on WT, Δ*mspA*, Δ*mspC*, the non-targeting sgRNA control (NT) and strains carrying either of two sgRNAs targeting *mspA* or *mspC*. KD, CRISPRi knockdown. Five successive phage dilutions are shown from left to right.

### MspA and MspC contribute to susceptibility in a phage-dependent manner

To determine whether the effects of *mspA* and *mspC* deletion were specific to BPsΔ33HTH HRM10 or extended to other mycobacteriophages, a panel of 25 phages representing multiple phage clusters and singleton phages was standardized to an initial titer of 10^10^ PFU/mL and tested on WT, Δ*mspA* and Δ*mspC* (Fig. 5A). Deletion of either porin altered susceptibility to a substantial proportion of the phages tested, although the magnitude of the phenotype was generally greater in Δ*mspA*. Eleven of the 25 phages showed a pronounced phenotype on Δ*mspA*, with titers reduced by more than 1-log_10_ or with no discrete plaques detected. Among the latter, Corndog and Marvin produced inhibition zones without discrete plaques, whereas D29 HRMGD40 and Dandelion produced neither plaques nor detectable inhibition zones. The effects of *mspC* deletion were generally less pronounced, although reduced plaque formation was also observed for several phages. In contrast, several phages showed little or no change following deletion of either porin. Giles displayed a distinct plaque morphology, producing larger and clearer plaques on Δ*mspA* than on WT despite only a modest change in relative titer (Fig. 5A,B). To further validate one of the strong phenotypes identified across the phage panel, D29 HRMGD40 was tested on the *mspA*- and *mspC*-targeting CRISPRi strains. Reduced plaque formation relative to the non-targeting control was also observed following CRISPRi-mediated repression of either gene (Fig. 4D). Together with the BPsΔ33HTH HRM10 validation experiments, these results support MspA as a host factor whose contribution to mycobacteriophage susceptibility is phage dependent.

**Fig. 5.**
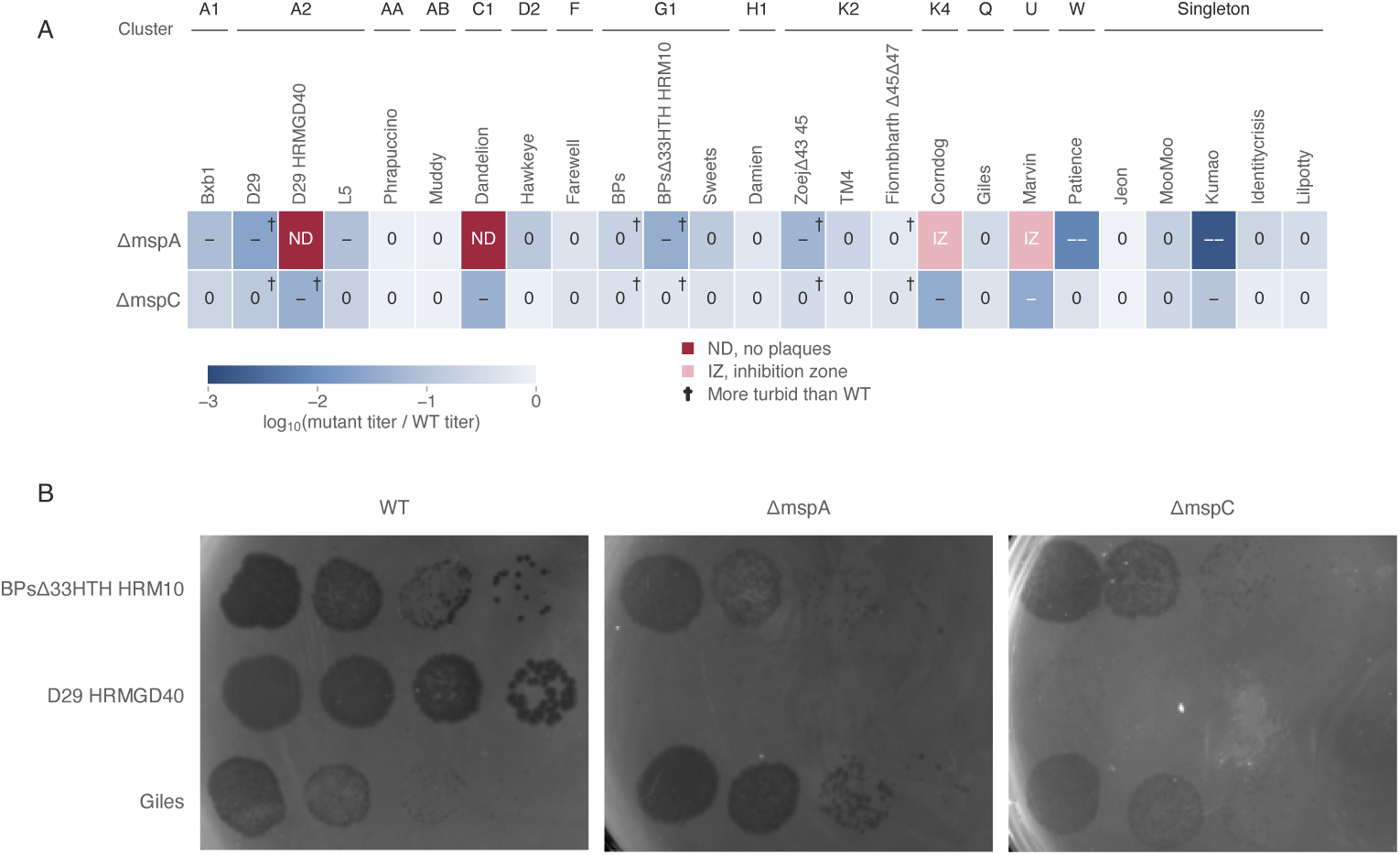
Phage-specific susceptibility profiles of *mspA* and *mspC* deletion mutants across mycobacteriophages. **A.** Titer change on Δ*mspA* and Δ*mspC* relative to WT for 25 mycobacteriophages, each applied at a standardized input titer of 10^10^ PFU/mL and grouped by cluster or subcluster; phages with no assigned cluster are grouped as singletons. Color, log_10_(mutant titer/WT titer). Cell labels: 0, <1 log_10_; *−*, 1 to <2 log_10_; *−−*, *≥*2 log_10_. ND, no detectable plaques; IZ, inhibition zone without discrete plaques; daggers, plaques more turbid than on WT. **B.** Spot assays for BPsΔ33HTH HRM10, D29 HRMGD40 and Giles on WT, Δ*mspA* and Δ*mspC M. smegmatis* mc^2^155, with phages by row and genotypes by column. Four successive phage dilutions are shown from left to right within each genotype.

## Discussion

The ability of bacteriophages to infect bacterial hosts depends on the recognition of specific surface components and the subsequent exploitation of cellular functions required for replication. In mycobacteria, most experimentally characterized determinants of phage susceptibility have been associated with cell envelope lipids and glycolipids. Trehalose polyphleates (TPPs), for example, are required for efficient adsorption and infection by the therapeutically relevant phages BPs and Muddy, while changes in surface lipid composition can impair infection by restricting access to these molecules. Although outer membrane proteins are well-established phage receptors in other bacterial species, their contribution to mycobacteriophage infection remains poorly understood. This is particularly relevant given the complex architecture of the mycobacterial envelope and the substantial variation in susceptibility observed among mycobacterial isolates. Identifying additional bacterial factors that influence infection is therefore important for understanding the molecular basis of mycobacteriophage host range. Here, we used genome-wide CRISPRi screening to identify bacterial functions associated with mycobacteriophage susceptibility, revealing the outer membrane porins MspA and MspC as previously unrecognized host factors. Independent genetic perturbations confirmed that loss or repression of either gene reduced susceptibility to BPsΔ33HTH HRM10, with a stronger phenotype following deletion of *mspA*. Interestingly, the closely related paralogs *mspB* and *mspD* were not enriched in the screen despite their high sequence similarity. MspA is the major outer membrane porin of *M. smegmatis*, and our proteomics measurements were consistent with a greater reduction in overall Msp-family abundance following its deletion. These observations suggest that the contributions of individual porins to phage infection may differ, although the relative importance of protein abundance and paralog-specific functions remains unresolved. More broadly, the identification of MspA and MspC extends the known determinants of mycobacteriophage susceptibility beyond the lipid components of the cell envelope. Importantly, the contribution of MspA was not conserved across the phages tested. Of the 25 mycobacteriophages examined, 11 showed a pronounced reduction in titer or loss of discrete plaque formation following *mspA* deletion, whereas the remaining phages were largely unaffected. The effect was not uniformly inhibitory, as Giles produced larger and clearer plaques in the absence of MspA. Particularly interesting was the comparison between parental phages and derivatives selected for expanded infectivity toward *M. abscessus*. Both D29 HRMGD40 and BPsΔ33HTH HRM10 showed greater sensitivity to MspA loss than their parental counterparts. These observations indicate that phages capable of infecting the same bacterial host can differ substantially in their dependence on individual envelope proteins. The differences between parental and host-range-adapted derivatives further suggest that changes associated with expanded infectivity can be accompanied by altered requirements for bacterial host factors. Whether these differences reflect changes in the recognition of MspA itself or in the interaction with other envelope components remains to be established. The precise role of MspA and MspC during infection remains unclear. BPsΔ33HTH HRM10 retained the ability to form plaques on both deletion strains, indicating that neither porin is individually essential for infection under the conditions tested. The porins could contribute directly to phage attachment, potentially alongside previously characterized surface determinants, or influence susceptibility indirectly by affecting envelope organization and physiology. MspA provides a major pathway for hydrophilic solutes across the mycobacterial outer membrane, and its loss can affect cellular permeability. Moreover, the high similarity and partially overlapping functions of Msp paralogs complicate the interpretation of individual deletion phenotypes. Our proteomics measurements detected residual Msp-family protein signal in both deletion strains but could not resolve the contribution of individual paralogs. Complementation and measurements of phage adsorption will therefore be important for determining whether the observed phenotypes arise from direct recognition of these proteins or secondary changes in the bacterial envelope. Beyond the Msp family, our CRISPRi screen also identified envelope-associated and tRNA-related functions whose repression altered bacterial fitness during phage exposure. Although these candidates remain to be validated, they highlight additional bacterial processes that may influence infection and demonstrate the broader utility of genome-wide CRISPRi for investigating mycobacteriophage host requirements. Understanding the bacterial factors that determine phage susceptibility is particularly important for therapeutic applications, where individual mycobacteriophages often infect only a limited subset of clinical isolates. Our findings establish that the protein composition of the mycobacterial outer membrane contributes to susceptibility in a phage-dependent manner and identify MspA as a host factor whose importance differs even between related phages. Extending these observations to clinically relevant mycobacteria will be necessary to determine whether natural variation in porin abundance or function contributes to isolate-specific susceptibility. Combining systematic genetic screening with experiments that resolve the molecular steps of infection could help establish how bacterial envelope composition and phage variation together determine infection outcomes, ultimately improving our ability to understand and predict mycobacteriophage host range.

## Materials and Methods

### Bacterial strains and phages

All bacterial strains and phages used in this study are listed in Supplementary Table S1. *M. smegmatis* and *M. abscessus* were cultured in Difco Middlebrook 7H9 broth supplemented with OADC and 1% arabinose, or on 7H10 agar supplemented with 10% glycerol and ADC, at 37 °C. When required, antibiotics were added at the following final concentrations: kanamycin (20 µg/mL), gentamicin (20 µg/mL), hygromycin B (50 µg/mL) and anhydrotetracycline (aTc, 100 ng/mL).

Phages were propagated on their bacterial hosts using a soft agar overlay method as previously described, with minor modifications[17, 18]. Briefly, 500 µL of mycobacterial culture was mixed with 50 µL of phage lysate and incubated at 37 °C for 1 h. The mixture was then combined with 5 mL of 7H9 medium containing 0.8% agar and overlaid onto a 7H10 agar plate. After incubation, the top agar layer containing phages was scraped off, vigorously vortexed, and centrifuged (4,000 *×* g, 10 min, 4 °C). The supernatant was collected and filtered through a 0.2 µm pore-size filter to remove residual bacteria. For titration, phage lysates were serially diluted in sodium chloride–magnesium sulfate (SM) buffer (100 mM NaCl, 10 mM MgSO_4_, and 50 mM Tris-HCl, pH 7.5) and spotted onto 0.8% top agar containing *M. smegmatis* mc^2^155 (final concentration, 1%).

### Bacterial challenge assay

Mid-exponentially growing *M. smegmatis* mc^2^155 cultures were infected with phages at the indicated multiplicities of infection (MOIs; 0.01, 0.1, 1, and 10) and incubated at 37 °C with agitation (200 rpm). Bacterial growth was monitored by measuring the optical density at 600 nm (OD_600_) at 30 min intervals using a microplate reader (SPECTROstar Nano, BMG Labtech, Germany).

### CRISPRi library transformation

The CRISPRi library RLC11 (#163955, Addgene) comprises 168,865 sgRNAs targeting 6,642 genes of *M. smegmatis* and 7,421 non-targeting sgRNAs serving as negative controls for normalization[15]. The library was amplified in *E. coli* ElectroMAX™ DH10B (18290015, Invitrogen) and sequenced for quality validation (see Genomic DNA extraction and library preparation for Illumina sequencing).

Electrocompetent cells of *M. smegmatis* were prepared as previously described[15, 19]. Briefly, mid-exponentially growing cultures were supplemented with glycine (final concentration 0.2 M) 3 h prior to harvesting. Cells were chilled on ice for 1.5 h, centrifuged (4,000 *×* g, 10 min, 4 °C), washed three times with ice-cold 10% glycerol, and resuspended in 1/100 of the original culture volume in ice-cold 10% glycerol.

For transformation, 200 µL of freshly prepared competent cells were mixed with 5 µg of RLC11 DNA, incubated on ice for 10 min, and transferred to a 0.2 cm electroporation cuvette. Electroporation was performed with a single pulse at 2.5 kV, 25 µF, and 1,000 Ω, after which cells were recovered in 5 mL of 7H9 broth at 37 °C for 6 h. Cells were then centrifuged (4,000 *×* g, 10 min, 4 °C) and plated on 7H10 agar plates containing kanamycin for selection. In parallel, 10 µL aliquots of recovered cultures were plated to assess transformation efficiency. This procedure was repeated 80 times, yielding a total of 1.1 *×* 10^8^ CFU, corresponding to ∼600-fold coverage of the 176,286 sgRNAs. After 3 days of incubation at 37 °C, colonies were scraped, pooled in 7H9 broth, and passed through a 10-µm pluriStrainer® (pluriSelect, #43-50010-03) to remove clumps. The declumped library was supplemented with glycerol (final concentration 15%) and stored at *−*80 °C for further use.

### Genomic DNA extraction and library preparation for Illumina sequencing

Mycobacterial cell pellets corresponding to 10 OD_600_ units were washed twice with PBS and used for genomic DNA extraction using the CTAB method[20]. Unless otherwise stated, all reagent concentrations are reported as final. Cell pellets were resuspended in GTE buffer (10 mM EDTA, 50 mM glucose, 25 mM Tris-Cl, pH 8) supplemented with lysozyme (1 mg/mL) and incubated overnight at 37 °C. Lysis was induced by addition of SDS (1%) and proteinase K (0.125 mg/mL) for 30 min at 65 °C, followed by treatment with NaCl (0.625 M) and CTAB (1%) for an additional 30 min at 65 °C. DNA was extracted with chloroform:isoamyl alcohol (24:1, v/v), and the aqueous phase was collected after centrifugation (16,000 *×* g, 10 min, 25 °C). Samples were treated with RNase A (50 µg/mL, 30 min, 37 °C) and sequentially extracted with phenol:chloroform:isoamyl alcohol (25:24:1, v/v) and chloroform:isoamyl alcohol (24:1, v/v). Genomic DNA was precipitated with sodium acetate (0.3 M) and two volumes of ice-cold 100% ethanol, incubated at *−*80 °C for 1 h, pelleted (16,000 *×* g, 10 min, 4 °C), washed twice with 70% ethanol, air-dried, and resuspended in elution buffer (10 mM Tris-HCl, pH 8.5) for downstream use.

The sgRNA region was amplified from genomic DNA using NEBNext Ultra II Q5 Master Mix with a pool of forward primers and unique indexed reverse primers as described previously[15]. PCR was performed according to the manufacturer’s instructions with the following cycling program: 98 °C for 45 s; 17 cycles of 98 °C for 10 s, 64 °C for 30 s, and 65 °C for 20 s; final extension at 65 °C for 5 min. Amplicons of ∼230 bp were size-verified by agarose gel electrophoresis, purified with AMPure XP beads using a 1.2*×* one-sided selection, quantified using a Qubit fluorometer (Thermo Fisher), and assessed for fragment size distribution with a Bioanalyzer (Agilent). Indexed PCR products were pooled at equimolar ratios to 10 nM, and 20% PhiX was spiked in to increase sequence diversity. Libraries were subjected to single-end sequencing on an Illumina NextSeq 2000 or NovaSeq X Plus platform (1 *×* 85 cycles and 6 cycles of i7 index).

### Pooled CRISPRi phage screening

After thawing, three 1 mL aliquots of *M. smegmatis* harboring the RLC11 library were each inoculated into 25 mL of 7H9 broth supplemented with kanamycin and incubated at 37 °C with shaking until stationary phase. Cultures were then diluted 1:50 into fresh 7H9 medium resulting in an OD_600_ of ∼0.02, and aTc was added to induce dCas9 expression and initiate sgRNA-mediated gene silencing. Following a 36 h pre-depletion, cultures were divided into two groups, with one treated with phages at an MOI of 1 and the other mock-treated with an equal volume of SM buffer. Both groups were incubated at 37 °C, and cell pellets equivalent to 10 OD_600_ units were collected at 18 h and 96 h by centrifugation (4,000 *×* g, 10 min, 4 °C) for subsequent genomic DNA extraction and sequencing.

### NGS data analysis

Sequencing reads were processed using cutadapt[21] with the linked adapter sequence GAGTACAAAAAC…TCCCA, corresponding to conserved sequences flanking the reverse-complemented sgRNA. Reads lacking the expected adapter structure were discarded. Trimmed reads were assigned to the reverse-complemented RLC11 sgRNA library using MAGeCK v0.5.9.5[22], generating counts for 176,286 sgRNAs. Library representation was assessed based on mapping efficiency, the proportion of zero-count sgRNAs, and the Gini index.

Gene-level enrichment and depletion were initially assessed using MAGeCK. Phage-treated samples were compared with no-phage controls, with three biological replicates per condition. Read counts were normalized using the non-targeting control sgRNAs, and gene-level log_2_ fold changes (L2FCs) were calculated using the alpha-median method. Genes were considered significant at FDR < 0.01 and |L2FC| *≥* 1, with at least two sgRNAs supporting the corresponding direction of change.

To assess time-dependent phage effects, genes identified as hits in the primary MAGeCK analysis were further analyzed using the MAGeCK maximum-likelihood estimation (MLE) framework[23]. Up to 20 sgRNAs per gene were retained based on their abundance in the 0 h no-phage samples, while all 7,421 non-targeting sgRNAs were retained for control-based normalization. Time and MOI 1 phage treatment effects were modeled jointly, and a reparameterized model was used to estimate the change in the phage-specific effect between 18 and 96 h. In this framework, the *β* score is the gene-level effect size estimated by the model and is interpreted on a log fold change scale: its sign indicates the direction of selection acting on a knockdown and its magnitude the strength of that selection, so *β* is used to assess both whether a gene is enriched or depleted and how strongly, allowing phage-associated effects to be compared across genes and between time points. Positive and negative *β* values indicated a shift toward relative enrichment and depletion at 96 h, respectively, and genes with FDR *≤* 0.05 and |*β*| *≥* 1 were considered to show significant time-dependent phage effects.

### Generation of target gene-deleted *Mycobacterium* spp

*M. smegmatis* mc^2^155 Δ*mspA* and Δ*mspC* mutants were generated by homologous recombineering using the Che9c gp60 and gp61 recombinase system as previously described[24], with minor modifications (Supplementary Fig. S1A). *M. smegmatis* mc^2^155 harboring pJV53 was cultured in Difco Middlebrook 7H9 broth supplemented with kanamycin (50 µg/mL) at 37 °C with shaking. Cultures were diluted to an OD_600_ of 0.03, grown to an OD_600_ of approximately 0.4, and induced with 2% acetamide for 4 h to express the Che9c recombinases. Cells were chilled on ice, harvested, washed with ice-cold 10% glycerol as previously described[25], resuspended in 1/100 of the original culture volume, and stored at *−*80 °C until use.

Allelic exchange substrates (AESs) for deletion of *mspA* (MSMEG_0965) and *mspC* (MSMEG_5483) were generated by PCR using Q5® High-Fidelity DNA Polymerase (New England Biolabs®, #M0493L). A 786 bp gentamicin-resistance cassette containing its native promoter was amplified from pPR2720[26] and assembled between approximately 0.63 kb upstream and downstream homology arms for each target gene (Supplementary Fig. S1B). The fragments were digested with HindIII and BamHI, ligated using T4 DNA ligase, and the assembled Δ*mspA*::*genta*^R^ and Δ*mspC* ::*genta*^R^ AESs were re-amplified by PCR and gel purified. Primer sequences used for AES generation and mutant validation are listed in Supplementary Table S2.

For homologous recombination, approximately 500 ng of purified AES was electroporated into 100 µL of competent *M. smegmatis* mc^2^155 pJV53 cells at 2.5 kV, 600 Ω and 10 µF. Cells were recovered in 900 µL of Middlebrook 7H9 broth for 4 h at 37 °C and plated on Middlebrook 7H10 agar containing kanamycin (50 µg/mL) and gentamicin (20 µg/mL). Plates were incubated for 5 days at 37 °C, and individual colonies were propagated in selective 7H9 broth before preparation of glycerol stocks.

Deletion of *mspA* and *mspC* was confirmed by PCR mapping of the corresponding chromosomal loci. Crude DNA templates were prepared by heating washed cell pellets at 95 °C for 20 min and removing cellular debris by centrifugation. PCRs were performed using primers positioned outside the homologous recombination regions together with primers specific for *mspA*, *mspC* or the *genta*^R^ cassette. Amplicons were analyzed by agarose gel electrophoresis, and replacement of each target gene with the *genta*^R^ cassette was confirmed from the resulting PCR profiles.

### Sample preparation for abundance proteomics

Whole-cell protein extracts were prepared from *M. smegmatis* mc^2^155 WT, Δ*mspA* and Δ*mspC* in four biological replicates per strain, grown as described under Bacterial strains and phages. Proteomic samples were prepared as previously described with minor modifications[27]. Briefly, 5 mL of bacterial culture was harvested by centrifugation at 3,220 *×* g for 10 min and washed once with PBS. Cell pellets were lysed in 8 M urea and 100 mM ammonium bicarbonate (ABC) containing protease inhibitors by bead beating with 0.1 mm zirconia-silicate beads at 6.5 m/s for two 45 s cycles. Lysates were clarified by centrifugation at 17,000 *×* g for 10 min, and protein concentrations were determined by BCA assay. For each sample, 100 µg of protein was reduced with 5 mM TCEP and alkylated with 10 mM chloroacetamide for 30 min each at room temperature, with alkylation performed in the dark. Urea was diluted to 1 M with 20 mM ABC, followed by overnight digestion at 37 °C with 1 µg sequencing-grade trypsin. Digestion was terminated by acidification with formic acid. Peptides were desalted using C18 columns and dried by vacuum centrifugation.

### LC–MS/MS acquisition

Dried peptides were resuspended in buffer A (0.1% formic acid), and 200 ng per sample were analyzed on a timsTOF Pro2 connected to a ProteoElute UHPLC (Bruker). Peptides were separated on a 150 mm *×* 150 µm C18 analytical column heated at 50 °C using a 90 min gradient at 0.5 µL/min. The capillary voltage was set to 1,600 V with a source temperature of 180 °C. Data were acquired in DIA-PASEF[28] with 48 asymmetric MS/MS isolation windows covering a precursor range of *m/z* 450–1250 and an ion mobility range of 0.85–1.30 1/*K* _0_. The window scheme was designed with py_diAID[29].

### Proteomics data analysis

DIA files were searched with DIA-NN (v1.8)[30] using a library-free approach against the *Mycolicibacterium smegmatis* mc^2^155 proteome (6,731 entries); no phage sequences were included. MS1 and MS2 mass tolerances were set to 5 and 10 ppm, respectively. Spectral library generation used precursor charge states 2–5, one missed cleavage and an MS1 mass range matching the DIA precursor range (*m/z* 450–1250). Protein grouping was performed on the library protein identifiers, and cross-run normalization was disabled. The global report was filtered to a protein group Q value of *≤* 1% (Lib.PG.Q.Value). Protein-group intensities were log_2_ transformed and median centered across samples. Missing values were not imputed, and a protein group was tested only when quantified in at least two replicates of both groups compared. Differential abundance between each deletion strain and WT was assessed using two-sided Welch’s *t*-tests on the normalized log_2_ intensities, and *p* values were adjusted across protein groups using the Benjamini–Hochberg procedure. Because the detected peptides do not distinguish MspA, MspB and MspC, these proteins are reported as a single shared protein group.

## Acknowledgements

We would like to thank Graham Hatfull (University of Pittsburgh, PA, USA) for providing the mycobacteriophages used in this study. We thank the members of the AF Lab for their valuable discussions and insights. AF and DK are supported by the SciLifeLab and Wallenberg Data-Driven Life Science Program (KAW 2020.0239) and the Swedish Research Council (2023-01970_VR). The funders had no role in study design, data collection and analysis, decision to publish or preparation of the manuscript.

## Author contributions

D.K.: Conceptualization, Investigation, Methodology, Validation, Visualization, Data curation, Writing – original draft. M.M.: Investigation, Methodology, Validation, Writing – original draft. K.A.E.: Methodology. M.A.D.: Methodology. G.I.P.: Investigation. A.G.N.: Investigation. D.R.P.: Investigation. P.S.: Resources, Methodology, Supervision. J.M.R.: Resources, Methodology, Supervision. A.F.: Resources, Funding Acquisition, Conceptualization, Methodology, Supervision, Writing – review & editing.

---

**Table S1.** Bacterial strains and bacteriophages used in this study.

| Strain | Relevant characteristics | Source/<br>reference |
| --- | --- | --- |
| *M. smegmatis* mc<sup>2</sup>155 (WT) | Wild type reference strain | ATCC, 700084 |
| *M. smegmatis* mc<sup>2</sup>155 $\Delta mspA$ | $\Delta mspA::genta^R$; pJV53 (Kan<sup>R</sup>) | This study |
| *M. smegmatis* mc<sup>2</sup>155 $\Delta mspC$ | $\Delta mspC::genta^R$; pJV53 (Kan<sup>R</sup>) | This study |
| *M. smegmatis* RLC11 pool | Pooled CRISPRi library used for genome-wide screening | Addgene, 163955 |
| *M. smegmatis* NT | CRISPRi strain carrying a non-targeting sgRNA | This study |
| *M. smegmatis* *mspA* KD, sgRNA 1 | Individual *mspA*-targeting CRISPRi strain | This study |
| *M. smegmatis* *mspA* KD, sgRNA 2 | Individual *mspA*-targeting CRISPRi strain | This study |
| *M. smegmatis* *mspC* KD, sgRNA 1 | Individual *mspC*-targeting CRISPRi strain | This study |
| *M. smegmatis* *mspC* KD, sgRNA 2 | Individual *mspC*-targeting CRISPRi strain | This study |
| *M. smegmatis* mc<sup>2</sup>155/pJV53 | pJV53 (Kan<sup>R</sup>); construction intermediate used to generate the deletion mutants | Addgene, 26094 |
| *E. coli* ElectroMAX DH10B | Host used for amplification of the RLC11 library | Invitrogen, 18290015 |

WT, wild type; NT, non-targeting; KD, knockdown; Kan<sup>R</sup>, kanamycin resistance; *genta*<sup>R</sup>, gentamicin-resistance marker.

**Table S2.**
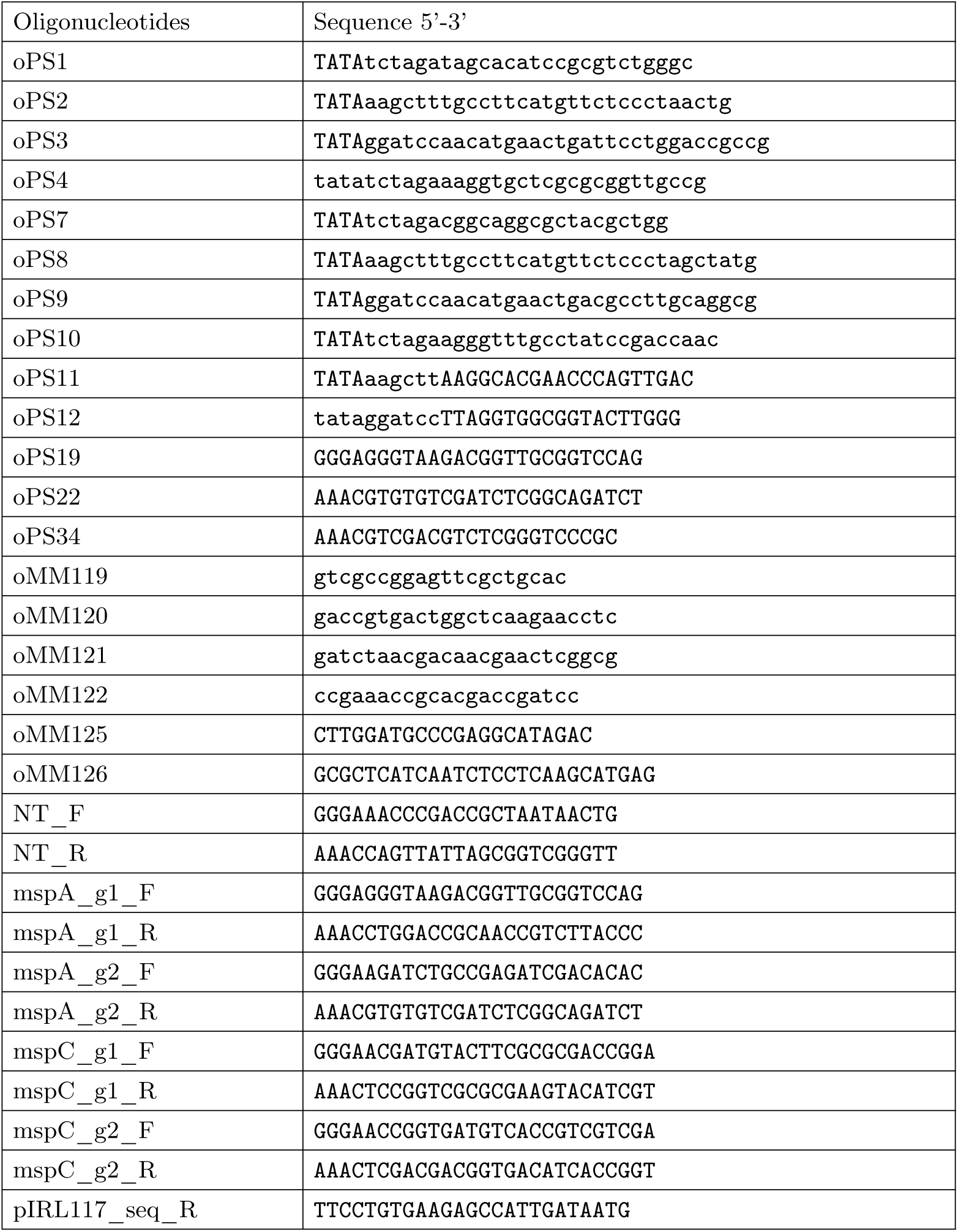
Oligonucleotides used in this study.

**Fig. S1.**
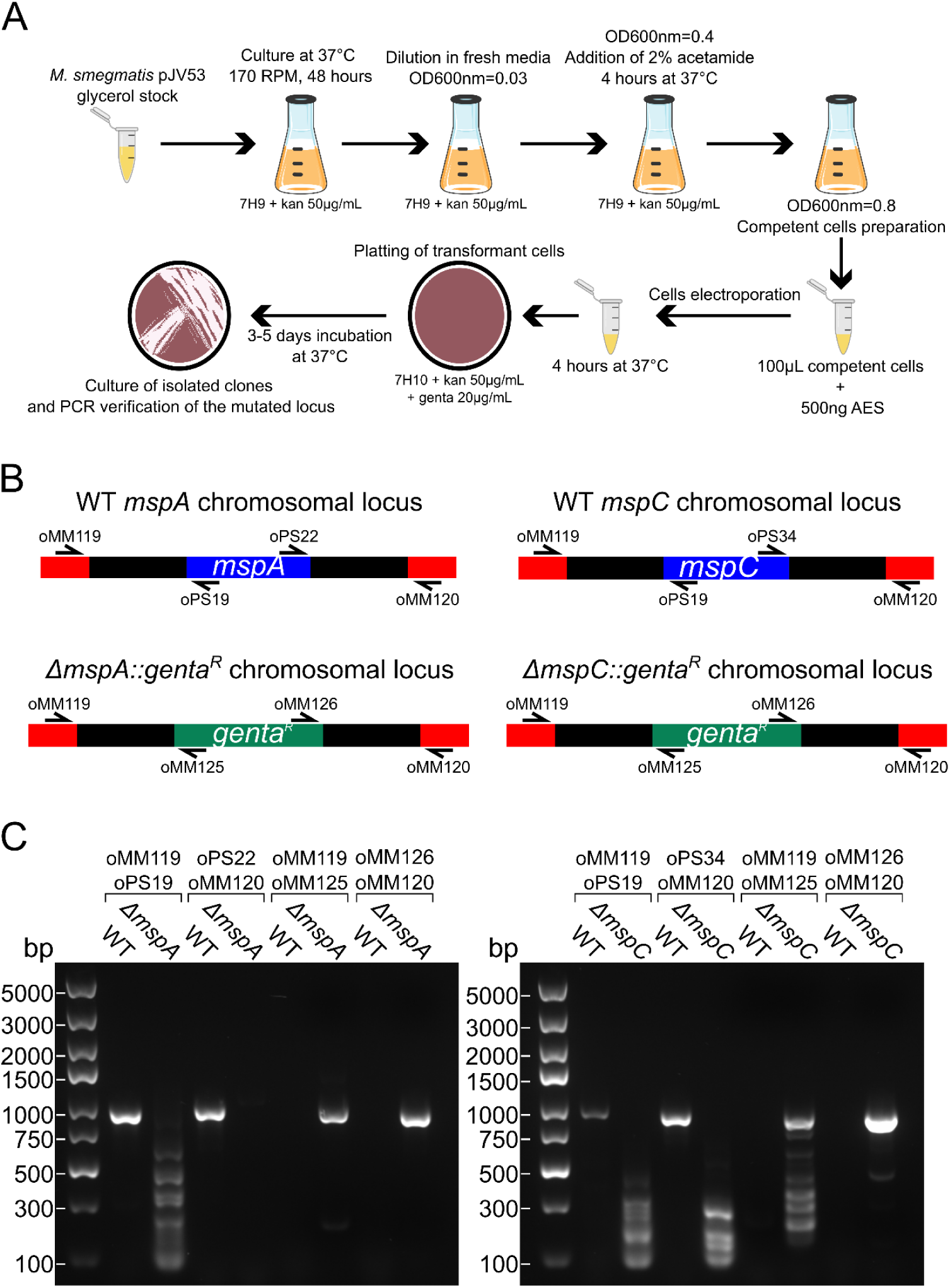
Generation and validation of *M. smegmatis* mc^2^155 Δ*mspA* and Δ*mspC* strains. **A.** Workflow used for homologous recombineering. *M. smegmatis* mc^2^155 carrying pJV53 was grown from a glycerol stock in 7H9 containing kanamycin (50 µg/mL) at 37 °C and 170 rpm for 48 h, diluted into fresh medium to an OD_600_ of 0.03, grown to an OD_600_ of 0.4, and induced with 2% acetamide for 4 h at 37 °C to express the Che9c recombinases. Competent cells were prepared at an OD_600_ of 0.8, and 100 µL were electroporated with 500 ng of allelic exchange substrate (AES). After 4 h of recovery at 37 °C, cells were plated on 7H10 containing kanamycin (50 µg/mL) and gentamicin (20 µg/mL) and incubated for 3–5 days at 37 °C. Isolated clones were then propagated and the modified locus verified by PCR. **B.** Maps of the *mspA* (left) and *mspC* (right) loci before (top) and after (bottom) replacement of the target gene with the *genta*^R^ cassette, together with the oligonucleotides used for PCR validation. Upstream and downstream homology arms are shown in red, the target gene in blue, and the *genta*^R^ cassette in green. Arrowheads indicate primer positions and orientations: oMM119 and oMM120 anneal outside the homology arms, oPS19 together with oPS22 (*mspA*) or oPS34 (*mspC*) anneal within the target gene, and oMM125 and oMM126 anneal within the *genta*^R^ cassette. **C.** PCR mapping of the *mspA* (left) and *mspC* (right) loci in WT and the corresponding deletion mutant. Four primer pairs were used per locus, each run as an adjacent WT/mutant pair: oMM119/oPS19 and oPS22/oMM120 (oPS34/oMM120 for *mspC*) amplify across the intact target gene, whereas oMM119/oMM125 and oMM126/oMM120 amplify across the left and right junctions of the *genta*^R^ replacement. Products were obtained from WT with the gene-specific pairs and from the deletion mutants with the cassette-specific pairs, and not from the reciprocal combinations. Amplicons were resolved by agarose gel electrophoresis alongside a DNA size standard (sizes in bp at left). Primer sequences are listed in Supplementary Table S2.

